# A computationally-enhanced hiCLIP atlas reveals Staufen1 RNA binding features and links 3’ UTR structure to RNA metabolism

**DOI:** 10.1101/2022.06.13.495933

**Authors:** Anob M. Chakrabarti, Ira A. Iosub, Flora C. Y. Lee, Jernej Ule, Nicholas M. Luscombe

## Abstract

The structure of mRNA molecules plays an important role in its interactions with *trans*-acting factors, notably RNA binding proteins (RBPs), thus contributing to the functional consequences of this interplay. However, current transcriptome-wide experimental methods to chart these interactions are limited by their poor sensitivity. Here we extend the hiCLIP atlas of duplexes bound by Staufen1 (STAU1) ∼10-fold, through careful consideration of experimental assumptions, and the development of bespoke computational methods which we apply to existing data. We present *Tosca*, a Nextflow computational pipeline for the processing, analysis and visualisation of proximity ligation sequencing data generally. We use our extended duplex atlas to discover insights into the RNA selectivity of STAU1, revealing the importance of structural symmetry and duplex-span-dependent nucleotide composition. Furthermore, we identify heterogeneity in the relationship between STAU1-bound 3’ UTRs and metabolism of the associated RNAs that we relate to RNA structure: transcripts with short-range proximal 3’ UTR duplexes have high degradation rates, but those with long-range duplexes have low rates. Overall, our work enables the integrative analysis of proximity ligation data delivering insights into specific features and effects of RBP-RNA structure interactions.

## INTRODUCTION

Interactions between RNA and associated trans-acting factors, notably RNA binding proteins (RBPs), are important for post-transcriptional regulation. It is becoming increasingly evident that the structure of RNA molecules plays an important role in this interplay. In particular, there are RBPs that contain protein domains that specifically bind double-stranded RNA (dsRNA) duplexes. The Staufen family is one such group of proteins with important roles in mRNA localisation, stability and translation. In order to understand the relationships between RNA structure, Staufen binding and the functional consequences, a comprehensive transcriptome-wide atlas of the bound structures *in vivo* is needed.

To this end, hiCLIP (hybrid individual-nucleotide resolution UV crosslinking and immunoprecipitation), an RNA proximity ligation method (Fig. 1A) was developed to study RNA duplexes bound by double-stranded RNA binding proteins (dsRBPs), such as Staufen1 (STAU1) (1, 2). UV-C is used to crosslink RBP-RNA complexes *in vivo*, following which cells are lysed and RNA partially fragmented by Rnase I. RBP-bound duplexes are enriched by immunoprecipitation, and the two fragments of interacting RNA strands ligated together. After digesting the bound RBP, cDNA is prepared from these molecules and sequenced. Two successfully ligated RNA fragments will yield a hybrid read, which we define as a sequencing read that maps non-contiguously to the transcriptome and thus represents two proximity ligated fragments (Fig. 1A, purple and green panels). Hybrid reads represent a duplex, which we define as a *unique* RNA structure formed by two strands of RNA - thus a duplex detected by hiCLIP can be supported by multiple hybrids. We refer to each hybrid or duplex as having two arms: the proximal arm is the 5’-most and the distal arm the 3’-most, each corresponding to one ligated RNA fragment. The non-ligated RNAs, which constitute the rest of the library, yield non-hybrid reads, which we define as a sequencing read that maps contiguously to the transcriptome and represents one RNA fragment. These reads may or may not represent a duplex (Fig. 1A, turquoise panel).

**Figure 1.**
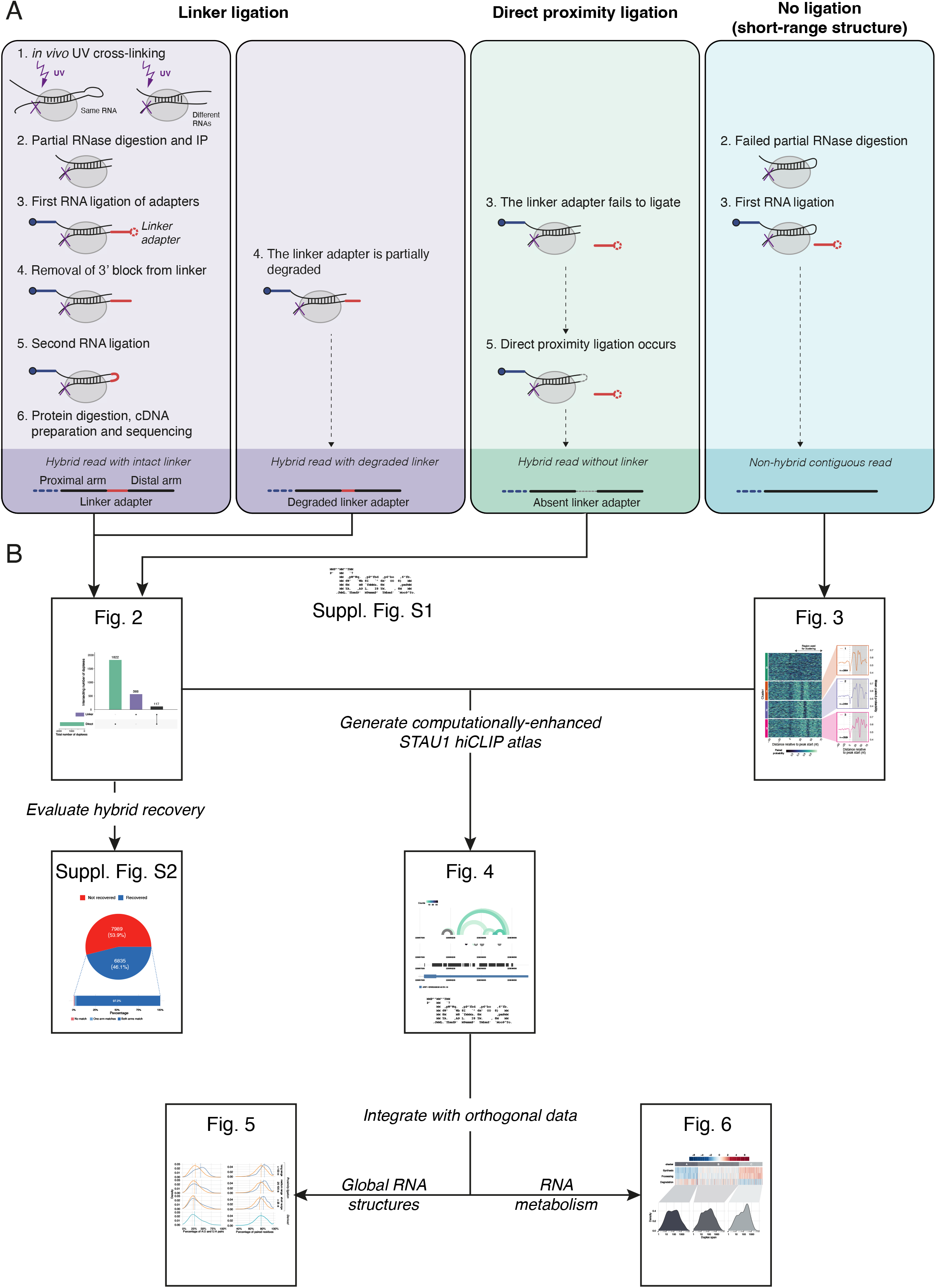
The hiCLIP method and sources of hybrid reads with a roadmap through the presented analysis and results. (A) Evaluation of the hiCLIP experimental steps to assess where in the data analysis duplexes could potentially have been “lost” in the original analysis (2) and the corresponding types of reads in each scenario. (B) A roadmap through the analysis, initially computationally-extending the atlas, then generating integrative insights, with the corresponding figures indicated.

Although proximity ligation experiments are primarily aimed at detecting hybrid reads, recovered hybrids typically constitute only a minority of the sequencing data due to both experimental and computational challenges: the ligation efficiency is low and accurate hybrid read identification and delineation of the original interacting RNA fragments is non-trivial. A data analysis challenge here is to determine unambiguously where the two arms start and end within sequencing reads, so that they can be mapped to the correct transcript locations. To address these issues, the hiCLIP method improved on previous methods (e.g. Crosslinking, Ligation And Sequencing of Hybrids; CLASH (3)), most notably by introducing a linker adapter to bridge the two RNA fragments into a single molecule instead of relying upon direct proximity ligation. This key innovation was devised to increase ligation efficiency (2) as well as streamline the computational workflow and ensure non-ambiguous assignment of the hybrid read arms to the correct transcriptomic loci. Therefore, the original computational pipeline searched for reads containing the linker adapter sequence flanked by the two arms. The two arms were individually mapped to the transcriptome, and then transcriptomic regions containing pairs of mapped arms originating from the same hybrid defined as duplex-forming. A drawback of this approach is that only 1-2% of all reads in the sequencing libraries were classified as hybrid (2). Ultimately, this meant that fewer than 1,000 duplexes in the transcriptome could be confidently identified (i.e. with more than 1 supporting hybrid read). Given the low sensitivity of duplex detection, we hypothesise that only a fraction of the *in vivo* STAU1 duplexes have been recovered from the data.

Moreover, the paucity of robustly-detected duplexes limits the extent to which hiCLIP results can be broadly integrated with other transcriptome-wide RNA duplex datasets, such as PARIS (4), or functional RNA metabolism datasets, such as 4sU-seq (5) to contextualise the Staufen-bound duplexes and gain further biological insights. While a role for STAU1 in RNA metabolism has been established for selected transcripts and regulatory mechanisms have been described, such as polysome association (6) and Staufen mediated decay (7, 8), a systematic transcriptome-wide exploration of the effect of STAU1-bound duplexes on RNA metabolism and stability has been lacking. Motivated to reveal the nature of STAU1 binding and its consequent effects on RNA metabolism on a transcriptome-wide scale, we aimed to improve hybrid detection from the STAU1 hiCLIP data - a prerequisite for more thorough investigations.

To improve the sensitivity of duplex detection from hiCLIP data, we refined and extended the computational approach. Specifically, we re-evaluated the experimental steps to understand where duplexes could potentially have been “lost” in the original data analysis through misassignation of non-hybrid reads. We challenged three key assumptions of specific steps of the experimental protocol through which this could have occurred (Fig. 1A).

First, we sought to recover hybrids containing truncated linker adapter sequences. 20-30% of the reads were found to contain linker-sequencing adapter dimers rather than sequencing adapters alone (2), suggesting that as the linker adapter is composed of ribonucleotides it is susceptible to degradation. Thus, this raises the possibility that a few nucleotides may have been degraded from the 3’ end of linker adapters, however these shorter linker adapters were originally not searched for in the hybrid read selection process.

Second, we assessed for the direct proximity ligation of duplex arms, resulting in hybrids lacking the linker adapter sequence. The original computational approach assumed that the vast majority of RNA hybrids contained the intermolecular linker due to the observed inefficiencies of circularisation and that sufficiently long stretches of single stranded RNA to enable circularisation would not remain after RNase digestion (2, 9). Thus hybrids arising from direct proximity ligation were previously not considered. However, we know from CLASH (3, 10) that intermolecular ligation can occur in many cases without a linker adapter. This has also been found in other subsequently developed proximity ligation methods (4, 11, 12). Hence, we hypothesised that there were a proportion of hiCLIP duplexes where the linker adapter failed to ligate (Fig. 1A, green step 3), but were still subjected to the second ligation reaction (Fig. 1A, step 5). In other words, direct proximity ligation of the two arms in the absence of a linker adapter was present in the hiCLIP data, provided the single-stranded overhangs were sufficiently long. However, such hybrid reads were assigned as non-hybrid in the original analysis as they would not contain the linker adapter.

Third, we identified short-range structures with undigested loops. RNase is used to digest the unprotected, primarily single-stranded RNA that connects the two duplex arms (Fig. 1A, step 2). However, short loops between the two arms in a stem-loop may be inefficiently cleaved, partly due to being sterically protected by the RBP from the action of the RNase. Although such stem-loops are isolated by virtue of being crosslinked to STAU1, sequencing reads arising from them will map to the genome as a contiguous sequence and thus be assigned as non-hybrids.

In summary, we predict that there will be three types of sequencing read containing a bound RNA duplex (Fig. 1A): i) hybrid read with a linker adapter, either full-length or truncated (purple); ii) hybrid read without a linker adapter containing a hybrid through direct proximity ligation (green); iii) non-hybrid read, containing a duplex with a short-range loop (turquoise).

Here, we evaluate each of these hypotheses (Fig. 1B) and in so doing present a Nextflow computational pipeline, *Tosca* (Suppl. Fig. S1), for the processing and analysis of proximity ligation experiments. In refining our understanding of the consequences of particular experimental steps and by computationally addressing their alternative outcomes, we obtain a ∼10-fold increase in identified duplexes from a previously published STAU1 hiCLIP dataset (Fig. 2, Suppl. Fig. S2 and Fig. 3). In addition we develop a computational approach to study local RNA structures around sites of RBP binding, which can be applied generally to any RBP of interest (Fig. 3). This extended set of STAU1 hiCLIP duplexes (Fig. 4) enables us to contextualise STAU1-bound duplexes in the universe of global RNA structures (Fig. 5) and furthermore, to derive insights into the relationship between STAU1 binding and RNA metabolism (Fig. 6).

**Figure 2.**
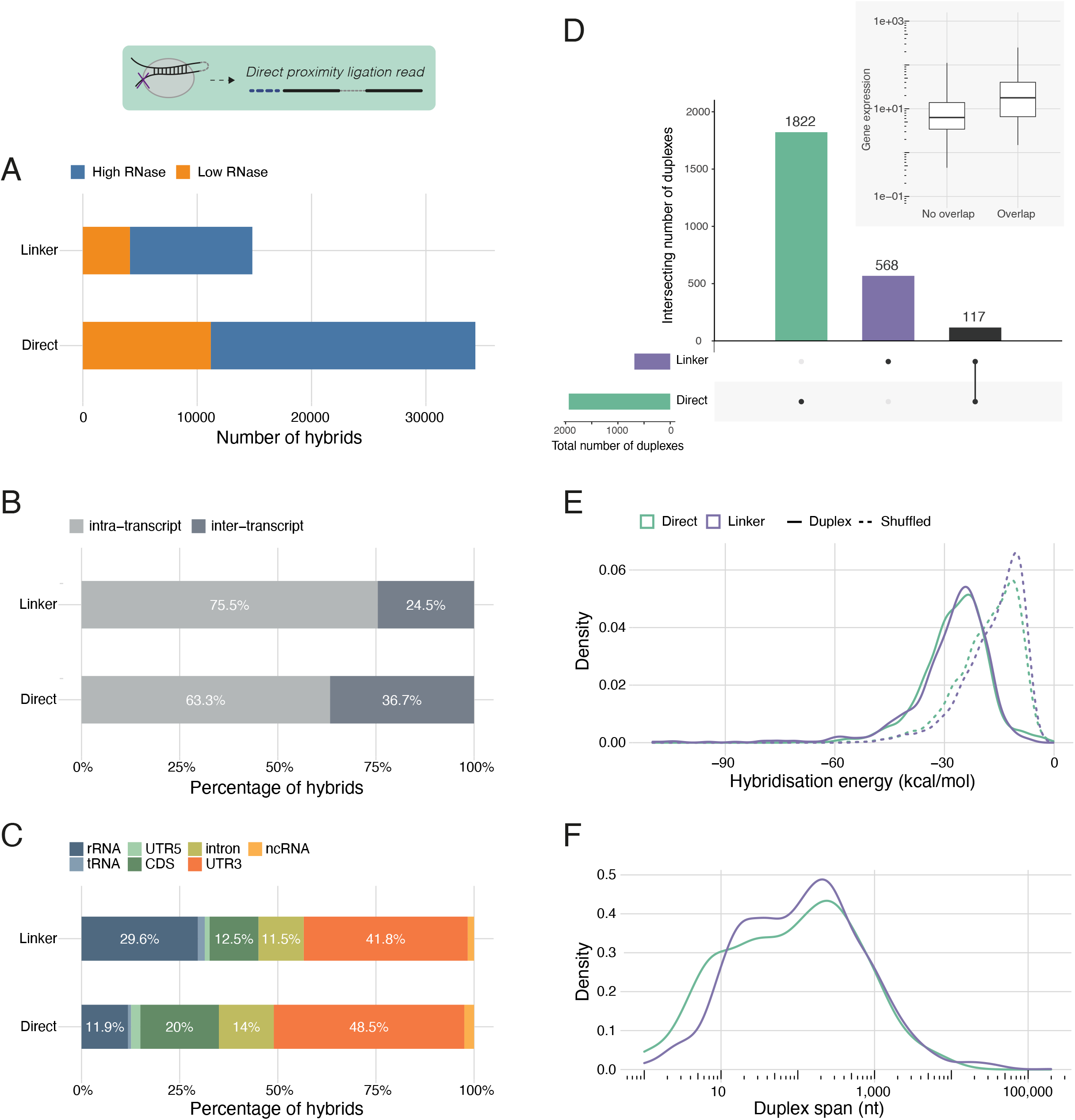
Direct proximity ligation is a major source of hybrid reads. (A) Comparison of hybrid counts across RNase concentration conditions for linker-containing hybrids and direct proximity ligation hybrids detected using *Tosca*. (B) Proportions of intra- and inter-transcript hybrids for the two hybrid read types (both RNase conditions pooled). (C) Regional distribution of hybrid arms for the two hybrid read types. (D) Overlap between confident duplexes derived from linker-containing hybrids and direct proximity ligation hybrids, respectively. Inset: gene expression of the genes on which duplexes were detected in both read types, compared with those in which duplexes were only detected in one. (E) Hybridisation energy of linker duplexes and direct proximity ligation duplexes compared to their corresponding shuffled controls. (F) Duplex spans (i.e. genomic distance between proximal and distal arms) of linker duplexes and direct proximity ligation duplexes.

**Figure 3.**
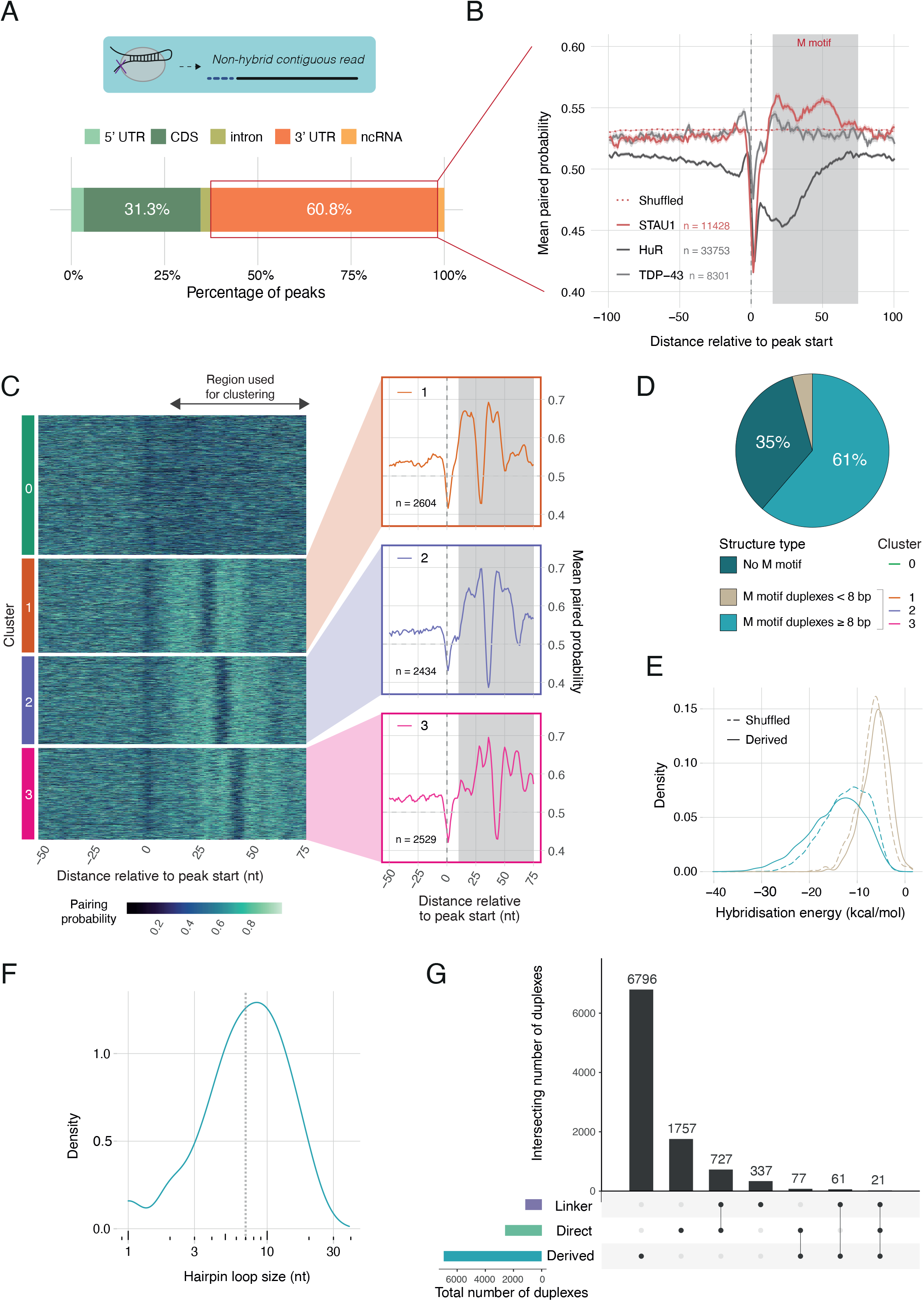
Deriving STAU1-bound short-range duplexes from crosslinking peaks. (A) Regional distribution of STAU1 peaks from non-hybrid reads in STAU1 hiCLIP data. (B) Paired probability metaprofiles centred on peak starts for three RBPs known to bind to 3’ UTRs (STAU1, HuR, TDP-43). For each RBP, the 3’ UTRs with peaks specific to each RBP were used to construct the metaprofiles (STAU1 - 11428 peaks, TDP-43 - 8301 peaks, HuR - 33753 peaks). The grey shaded area (+10…+75 nt) indicates the relative area spanned by the “M” shaped profile specific to STAU1. (C) Heatmap of clustered paired probability profiles for STAU1 3’ UTR peaks. The clusters were generated by k-means clustering using the +10…+75 nt region relative to peak starts. On the right, the corresponding mean probability profiles for the three clusters with high paired probability downstream STAU1 (i.e. occurrence of “M” shaped profile). (D) Proportion of peaks with or without predicted downstream duplexes and their classification based on the number of paired residues. (E) Hybridisation energy of the derived short-range duplexes compared to corresponding shuffled controls. (F) Hairpin loop size distribution for STAU1 derived stem-loop duplexes. (G) The final enhanced atlas of STAU1 hiCLIP duplexes comprising the full set of hybrids from proximity ligation (direct proximity ligation or via the linker) and 3’ UTR derived structures. The number of duplexes supported by each or multiple sources is indicated.

**Figure 4.**
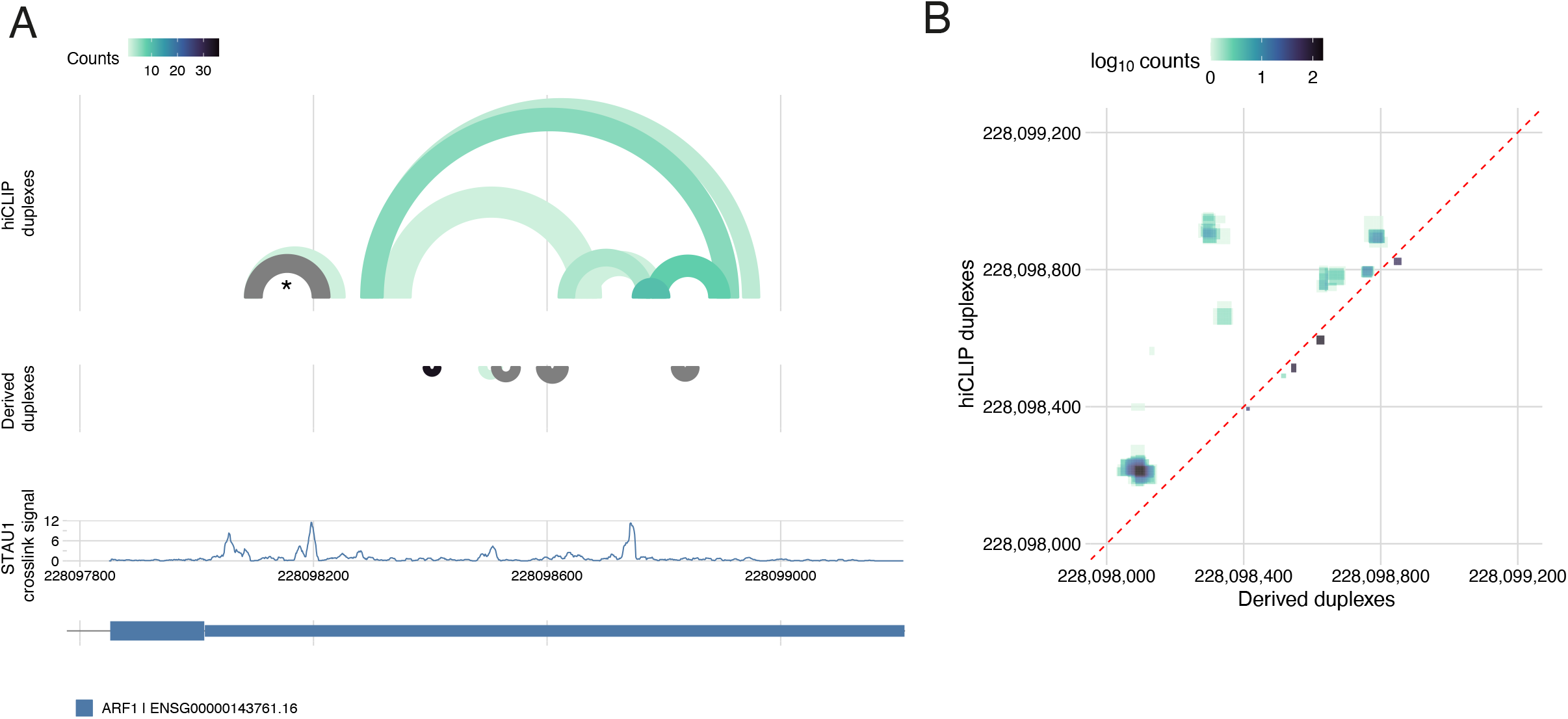
*Tosca* incorporates visualisation capabilities to integrate identified duplexes with other genomic datasets. (A) Arc plot of the experimental hiCLIP range and short-range derived duplexes on the *ARF1* 3’ UTR, alongside STAU1 crosslinking signal and transcriptomic annotation; the known structure re-discovered in the STAU1 hiCLIP data is marked with an asterisk. (B) Contact map matrix showing counts at each base pairing location with experimental hiCLIP duplexes above the diagonal (dashed red line) and derived duplexes below.

**Figure 5.**
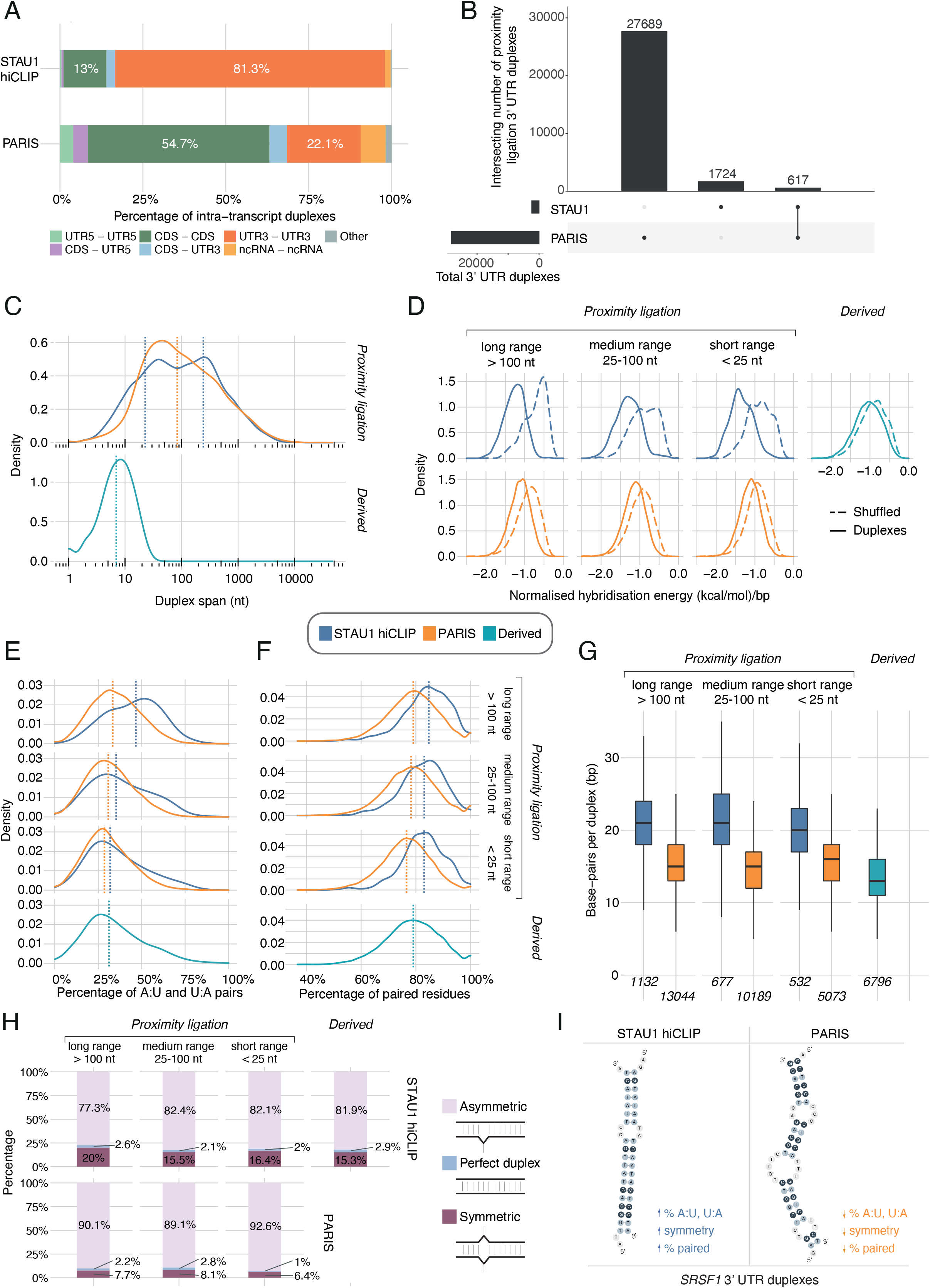
Contextualisation of STAU1 hiCLIP duplexes in the global universe of duplexes detected by PARIS highlights features driving STAU1 RNA selectivity. (A) Regional distribution of intra-transcript pairwise interactions between proximity ligation STAU1 hiCLIP and PARIS duplexes. (rRNA, tRNA and intronic duplexes were excluded from this analysis.) (B) Overlap between STAU1 hiCLIP and PARIS 3’ UTR intra-transcript proximity ligation duplexes. (C) Duplex spans of 3’ UTR intra-transcript duplexes. Vertical dashed lines indicate the median span for each detection method. For the bimodal distribution (STAU1 hiCLIP proximity ligation duplexes) the median of each normal distribution defined by the Gaussian mixture model applied in Suppl. Fig. 5 A is indicated. (D) Normalised hybridisation energy of 3’UTR intra-transcript duplexes compared to shuffled controls. (E) A:U and U:A base-pair percentage distributions. (F) Percentage of paired residues within the duplexes. (G) Number of paired residues within the duplexes. (H) Classification of 3’ UTR duplexes based on i) the absence of bulges (perfect duplex) or ii) the relative positioning and sizes of the bulges in the proximal and distal arms (symmetric and asymmetric). Proportions of 3’ UTR duplex types in STAU1 hiCLIP and PARIS data, respectively, grouped according to duplex span. (I) Summary of the STAU1 hiCLIP selectivity features (nucleotide composition and base-pairing architecture) compared to PARIS for example duplexes on the *SRSF1* 3’ UTR (see also Suppl. Fig. S5E). Distributions in (D-F) were compared with the Mann-Whitney test.

**Figure 6:**
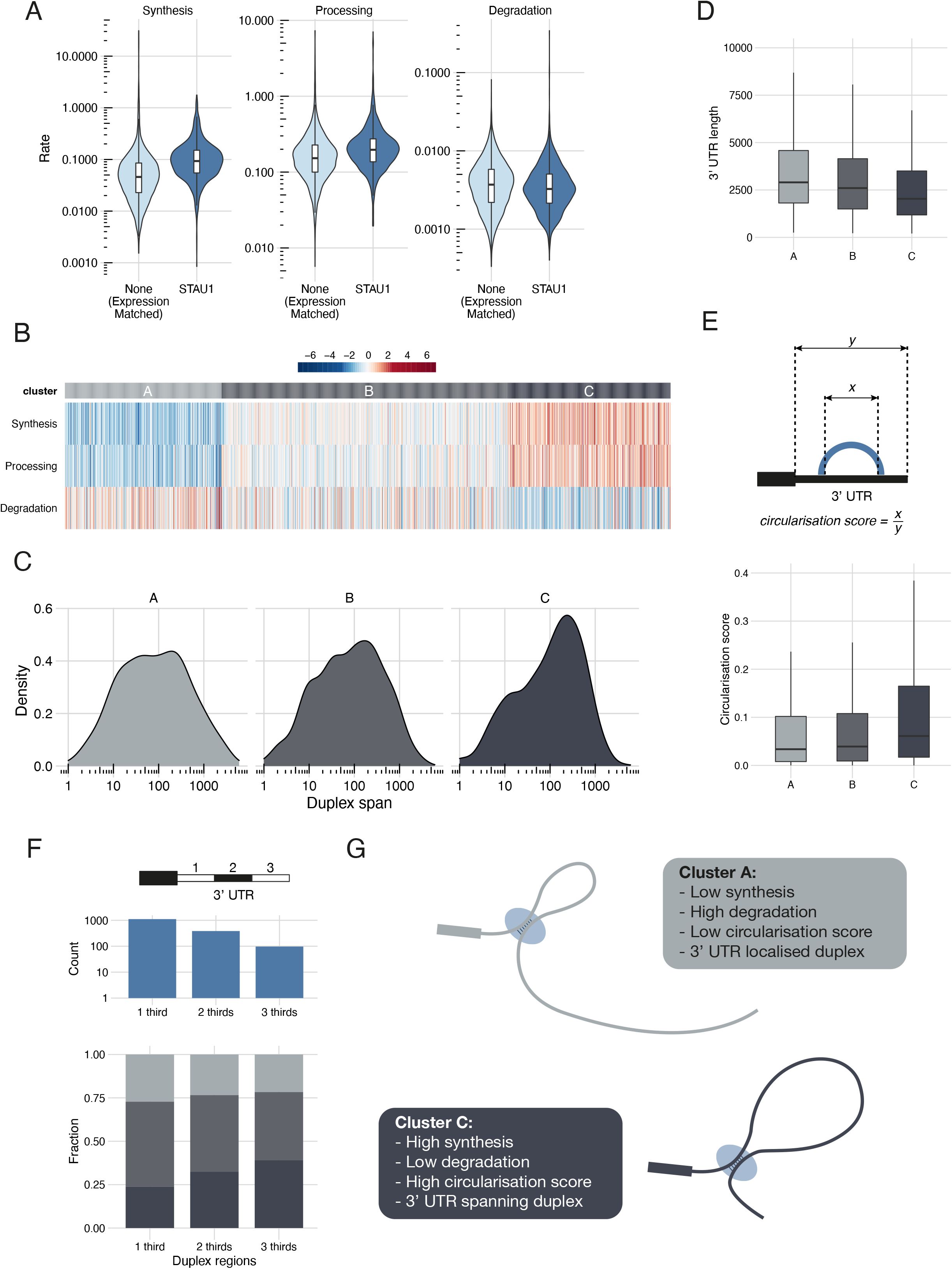
The relationship between STAU1-bound 3’ UTR duplexes and RNA metabolism. (A) RNA metabolism (synthesis, processing and degradation rates) as measured by (5) for genes with 3’ UTR duplexes identified by proximity ligation in STAU1 hiCLIP compared to genes without duplexes but matched for gene expression. (B) RNA metabolism profiles cluster based on synthesis, processing and degradation rates estimated by (5) for genes with STAU1-bound duplexes. (C) Mean 3’ UTR duplex spans per gene for each cluster in (C). (D) 3’ UTR lengths of the genes in each cluster. (E) Above: schematic describing the circularisation score calculation. Below: circularisation scores for the genes in each cluster. (F) Above: Number of genes with duplexes spanning one-, two- or three-thirds of the length of the 3’ UTR. Below: Distribution of metabolism clusters for genes with duplexes spanning the three sub-segments of the 3’ UTR (G) Schematic model indicating the two types of 3’ UTR duplex bound by STAU1 and their relationship with RNA degradation.

## MATERIAL AND METHODS

### Data provenance

Raw Staufen1 hiCLIP sequencing data from (2) was downloaded from ArrayExpress (E-MTAB-2937) and demultiplexed into the two RNase conditions (high and low). Three replicates of raw PARIS sequencing data in HEK293T cells were obtained from GEO (SRR2814763-5). For all experiments, each replicate was processed separately as described below.

### Reference annotation and sequence creation

For all analyses we used the GRCh38 build of the human genome with the Gencode V33 annotation. We created a custom reference sequence for the alignment of hybrid reads. This was necessary to reduce the complexity of the alignment problem for direct proximity ligation reads (i.e. without a linker adapter) to make it computationally tractable; largely by removing unannotated regions of the genome. We started to create this reference sequence by combining rDNA and 5S rRNA sequences from NCBI (U13369.1 and NR_023363.1); mature tRNA sequences from gtRNAdb (13). We then selected genes annotated in Gencode V33 as: “protein_coding”, “IG_[A-Z]_gene” and “TR_[A-Z]_gene” as protein coding genes and those annotated as “lncRNA” or “vault_RNA” as non-coding. We collapsed overlapping genes, concatenating the gene names, to avoid duplicating the same region in our reference sequence.

To obtain the corresponding sequences, first we created a mask that contained the genomic coordinates of regions: (i) annotated in Gencode V33 as: “rRNA”, “rRNA_pseudogene”, or “snRNA”; (ii) annotated in gtRNAdb as tRNA; and (iii) annotated by RepeatMasker (obtained from the UCSC table browser) as: “rRNA”, “tRNA”, “snRNA”, “srpRNA”, “scRNA”, or “RNA” (the latter corresponding to 7SK). We used this as input to BEDtools (14) to mask the GRCh38 sequence before obtaining the sequences for each of our selected RNAs to create our reference.

### Identification and alignment of hybrids with a linker adapter

To identify duplexes from hybrid reads with a linker present we broadly followed the originally published method (1, 2). As previously, we used Cutadapt (15) to trim the 3’ sequencing adapters and linker and sequencing adapter concatemers. Then we identified reads containing the full-length linker adapter. We additionally examined the reads for truncated linker adapters missing either the last or the last two nucleotides. We kept reads for which there were at least 12 nt sequences flanking the linker adapter, splitting out the two as the two hybrid arms for alignment.

As it was not possible to perform the original iterative step-wise alignment (first to rRNA, then tRNA, then mRNA and ncRNA) for direct proximity ligation reads without a linker adapter, in order to ensure consistency we also aligned each hybrid arm in one step to our reference sequences using STAR v 2.7.7a (16).

To mimic direct proximity ligation (i.e. hybrid reads without a linker adapter) *in silico* for these reads which we knew contained hybrids, we stitched together the two hybrid arm sequences after removing the intervening linker adapter sequence.

### *Tosca*: a Nextflow pipeline for proximity ligation data analysis and visualisation

To analyse hybrid reads created through direct proximity ligation i.e. from sequencing reads without a linker adapter, we developed a computational analysis pipeline, *Tosca*, in Nextflow (17) (Suppl. Fig. S1). *Tosca* first performs hybrid read identification, alignment and annotation; and duplex delineation using a graph-based clustering of hybrid read alignments, hybridisation energy calculation and generation of IGV tracks and plots for visualisation. The default parameter settings detailed below can all be optionally customised.

#### Identification and alignment of hybrids created through direct proximity ligation

Our proximity ligation hybrid read identification approach is inspired by the hyb pipeline (10) originally written for CLASH data. The basis of the method is to derive an optimal solution to the problem that a given sequencing read contains two sections that each align to two different parts of the reference, but that the location of the join between the two in the read sequence is unknown. As input to this stage of the analysis we use all reads in which a linker adapter (full-length or truncated) had not been detected above. First we use pblat (18), using a step size of 5, tile size of 11 and minimum score of 15 (10) to identify all partial read alignments to the reference sequence. Next we filter the pblat output using a stringent e-value threshold of ≤ 0.001 to keep only high quality alignments. We also filter out reads with more than 100 partial alignments to keep the next step computationally tractable. Then we select all the best partial alignments for a given set of read start and end coordinates. If a read contains one partial alignment that spans the majority of the read, leaving only 15 nt unaligned, then it is deemed not to contain a hybrid and is filtered out. This prioritises non-hybrid solutions over hybrid solutions and helps control the false-positive detection of hybrids. If the read is not filtered out, then all the best partial alignments are cross-joined with each other to derive all possible combinations. These combinations then pass through three filters. First, any partial alignments that have more than 4 nt overlap or are more than 4 nt apart in the read sequence are removed. A small amount of leeway is allowed as feasibly a few nucleotides at the ends could be assigned to either hybrid arm. Second, any solutions where both partial alignments align to the same reference sequence, but that overlap regions of the reference sequence are removed. Third, those solutions that start more than 5 nt away from the crosslink position are removed. At this point, for reads that have a unique solution, this is selected as the hybrid alignment. For reads that have more than one solution, any solution that overlaps one of the unique solutions is selected as the hybrid alignment. If there is more than one overlap, then the solution with greatest total length of the read aligned is selected. Any reads that still have multiple potential solutions are deemed ambiguous and not analysed further as hybrid reads. PCR duplicates are then collapsed using the unique molecular identifiers with the directional method from UMI-tools (19) modified to consider the RNA transcript and start coordinates of both arms of the aligned hybrid.

#### Delineation of duplexes using graph-based clustering of hybrids

To delineate duplexes we developed a graph-based clustering approach to overcome limitations of the original coverage-based approach when dealing with larger datasets with greater complexity. We used BEDtools (14) to identify overlapping hybrids (requiring both hybrid arms to overlap) and then calculated the fraction of the overlap for each arm as a proportion of the total span of the two overlapping arms. Those with a fraction ≥ 0.5 for both arms were annotated as valid overlapping hybrids. Then we used these calculations to create an undirected graph using the igraph package (20) where nodes were hybrids and edges overlaps (weighted by the fraction overlap). From this graph, we identified the connected components (i.e. set of linked vertices) to derive subgraphs of hybrids that represented the same duplex that we termed clusters. To define the ends of a duplex from the clustered hybrids, we either took the median or the maximum.

#### Calculation of hybridisation energy

To calculate the hybridisation energy for each duplex we used RNAduplex from the ViennaRNA package (21) to predict the minimum free energy duplex structure from the two arms, disallowing lonely pairs. We also calculated a control hybridisation energy for each duplex as the mean minimum free energy from 100 iterations of shuffling the duplex arm sequences (preserving dinucleotide content) using uShuffle (22).

#### Visualisation of hybrids

Comprehensive hybrid data are stored as tables and can be used to visualise hybrids or duplexes as arc plots. Additionally, the relevant attributes from the hybrid tables are also used to generate BAM files to enable visualisation in genome browsers. Custom ‘ID’, ‘XP’, ‘CL’, ‘RO’, and ‘BE’ BAM tags are used to enable grouping or colouring by hybrid id, experiment or sample, duplex cluster, hybrid read orientation and hybridisation energy respectively in, for example, IGV (Integrative Genomics Viewer). Furthermore, duplexes are converted into BED files for visualisation. Optionally, selected genes of interest can be supplied and arc plots generated for viewing in IGV and nucleotide-resolution or binned contact map matrices for static plotting.

### Identification of RBP crosslink peaks from non-hybrid hiCLIP and iCLIP

To identify STAU1 crosslink peaks from hiCLIP reads in which no hybrids had been detected, and HuR and TDP-43 crosslink peaks from iCLIP data (E-MTAB-11854 and (23) respectively) we used the nf-core/clipseq (v. 1.0.0 - Ianthine Pelican) pipeline with default parameters. Biological replicates for each RBP were processed separately to identify crosslinks and then merged using BEDTools (14) to create the input for the iCount peak caller. Significant crosslink sites were identified using a half-window setting of 10 nt for STAU1 and TDP-43 (on account of the broader binding profiles of these RBPs) and the default 3 nt for HuR. Peaks were calculated by merging significant crosslink sites within 10 nt (for STAU1 and TDP-43) and 3 nt (for HuR) of each other.

### Prediction of bound short-range duplexes from STAU1 non-hybrid reads

#### Derivation of pairing probability profiles

For all genes bound by STAU1, we defined a representative 3’ UTR. This was selected by processing the matched HEK293 RNA-seq data from (2) using the nf-core/rnaseq pipeline (v3.1 - Lead Alligator) with pseudoalignment using Salmon (24) to obtain transcripts per million values for each transcript. The 3’ UTR from the most highly expressed transcript for each bound gene was selected. If transcripts for a bound gene were not detected in the RNA-seq (likely due to low abundance), we selected the longest annotated transcript isoform (with ties broken by a hierarchy of most exons followed by longest 3’ UTR). For each 3’ UTR sequence, we calculated the local pairing probability for each base using RNAplfold from ViennaRNA (21) using a window size of 100 nt. We also generated control pairing probabilities as the mean of the pairing probability from 100 iterations of shuffling the 3’ UTR sequence (preserving dinucleotide content) using uShuffle (22).

We identified the probabilities in the −100 to +100 nt region centered on the STAU1 peak starts and for the metaprofiles calculated the mean and standard error of the mean. We performed the same analysis for HuR and TDP-43. To cluster the STAU1 peaks based on the downstream structure profile, we extracted the probabilities in the +10 to +75 nt region and performed k-means clustering. We applied the silhouette method to guide the choice of number of clusters.

#### Derivation of predicted STAU1-bound duplex structures

To predict duplex structures downstream of STAU1 crosslink sites, we selected the peaks from the clusters with evidence of a downstream stem-loop structure. From the metaprofiles for each cluster we calculated the regions that contained the proximal and distal duplex arms relative to the peak start position. To identify the boundaries of these regions we calculated the local minima of the profiles, which described the arm ends. For each peak, we then extracted the sequences for these regions and used them as input to RNAduplex to identify the duplex contained within and trimmed unpaired flanking nucleotides.

### PARIS data processing

We trimmed sequencing adapters using Cutadapt and collapsed PCR duplicates in advance of alignment using the readCollapse.pl script used in the original publication (available at https://github.com/qczhang/icSHAPE). Subsequent processing was using *Tosca* as above.

### Integration with RNA metabolism data

We used the processed data available from (5) quantifying RNA abundance (gene copy number) and RNA metabolism rates (synthesis, processing and degradation). To cluster the genes using the three rates, we log_10_-transformed the data and scaled and centred synthesis, processing and degradation rates to give them equal weighting in the clustering. We then used k-medoid clustering with 3 clusters, as identified using the silhouette method.

To identify relationships between STAU1-bound duplex span and 3’ UTR length, we first identified a representative 3’ UTR for a gene containing a duplex by selecting the longest 3’ UTR from the gene that overlapped the duplex. To correct duplex span and 3’ UTR length for intronic length (given that the STAU1 hiCLIP performed in (2) is cytoplasmic and thus reflects mRNAs), we identified 3’ UTR introns contained within either the duplex span or the 3’UTR and subtracted them respectively. We then calculated the circularisation score as the duplex span divided by the 3’ UTR length. This is analogous to the circularisation score defined previously (11), but substituting 3’ UTR length for gene length.

## RESULTS AND DISCUSSION

### Hybrid reads contain intact and degraded linker sequences

We first reprocessed the publicly available STAU1 hiCLIP data (Sugimoto et al. 2015; Materials and Methods) by broadly replicating the original analysis. The dataset consists of two replicates (high and low RNase library preparation conditions) and contain 2,429,385 and 2,996,034 reads respectively. One of the original rationales for the linker adapter was to enable easy bioinformatic deconvolution of the two hybrid arms from a sequencing read (2). We identified the location of the full linker sequence in the read (1) to extract 62,906 and 43,982 reads respectively, giving a total of 106,888 reads that contained hybrids. This compared with 72,280 and 46,502 in the original analysis, the difference owing to more stringent criteria (2).

First, we sought to recover hybrids containing degraded linker adapters (Fig. 1A, purple). We therefore also examined the sequenced reads for truncated linker adapter sequences (with up to 2 nt lost from 3’ end), and recovered a further 8,324 and 5,412 sequencing reads from the high and low RNase conditions respectively. Hence, although there is some degradation of the linker adapter, this is not a major source of lost hybrids.

Combining these two groups of linker hybrids, it was possible to reconstruct both arms after alignment for 20,091 and 12,184 hybrids in the high and low RNase conditions respectively. This compares with 21,291 and 14,067 in the original, where the more sensitive iterative mapping approach was able to be used. After UMI-aware collapsing of PCR duplicates, this left 10,705 and 4,119 unique linker hybrids. Overall, this meant that 2,329,526 (95.9%) and 2,894,708 (96.7%) sequencing reads were classified as not containing linker adapters in the high and low RNase conditions respectively and formed the starting point for the subsequent analysis.

### Hybrid reads can form through direct proximity ligation in the absence of the linker adapter

Next, we explored the occurrence of hybrids from direct proximity ligation. This challenged the validity of the assumption that a linker adapter was necessary for proximity ligation. However, identifying the two arms of a hybrid within a sequencing read without the benefit of the linker to demarcate the join is challenging. Using a partial alignment and hybrid reconstruction approach on a read-by-read basis (implemented in the *Tosca* computational pipeline, Suppl. Fig. S1, see Materials and Methods), we identified 92,422 and 58,186 such direct proximity ligation hybrid reads in the high and low RNase conditions respectively. Of these, 17,434 and 12,144 had more than one possible hybrid solution and so were filtered out as ambiguous. After UMI-aware collapsing of PCR duplicates, this left 23,129 and 11,195 unique direct proximity hybrids. This was over twice as many as were detected with a linker (Fig. 2A). There were similar proportions for both low and high RNase conditions, suggesting that RNase digestion within this range does not preclude proximity ligation. As for hybrids with a linker, the majority of direct proximity ligated hybrids were intra-transcript (linker, 75.5% and direct, 63.3%, Fig. 2B). While there were proportionally fewer proximity ligation hybrids recovered from rRNA regions compared to hybrids with a linker (linker, 29.6% and direct, 11.9%, Fig. 2C) - likely due to reaching multimapping limits or filtering out of ambiguous solutions - within mRNAs 3’ UTR binding dominated to a similar level (linker, 41.5% and direct, 48.5%, Fig. 2C).

We then used the hybrid reads from each source to identify the duplexes they represented using a graph-based clustering approach. Using solely linker hybrids identified 685 mRNA duplexes. However, including the direct proximity ligation hybrids increased the yield ∼4-fold to 2,507 (Fig. 2D). Interestingly, only a minority - 117 - of all duplexes were found from both sources (Fig. 2D). We found that genes with duplexes found in both datasets had a higher RNA abundance (Fig. 2D, inset). Thus, the relatively low overlap likely reflects the sensitivity limits of this hiCLIP dataset. As we are sampling from the population of STAU1-bound duplexes, those on more abundant transcripts are more likely to be recovered by both approaches. Next, we focused on intra-transcript duplexes in mRNA and ncRNA: this showed that duplexes identified from direct proximity ligation reads had similar hybridisation energies (Fig. 2E) and duplex spans, i.e. genomic distance between the two duplex arms (Fig. 2F) to the ones identified from linker hybrids. So, the features of hybrids detected by linker and direct proximity ligation approaches are highly comparable, demonstrating that the direct proximity hybrids also originate from STAU1-bound duplexes. Thus, the linker and direct proximity ligation hybrids were combined for our subsequent analysis.

### *In silico* read reconstruction shows not all hybrids are recoverable without a linker adapter

Next, we wanted to assess the limitations of computational hybrid recovery of direct proximity ligation hybrids. The STAU1 hiCLIP dataset is uniquely placed for evaluating the performance of hybrid detection approaches because the linker unambiguously delineates each hybrid arm and their exact transcriptomic locations - thus, the linker hybrids constitute a “ground truth” set of hybrids. To compare *Tosca* hybrid solutions to those identified by individual arm mapping, we took all the hybrid reads that contained a linker and removed the intervening linker adapter sequence. In this way we created 14,824 *in silico* direct proximity ligation hybrids, but for which we knew reliably both that they were hybrid reads and the transcripts and coordinates of the hybrid arms.

Processing these *in silico* hybrid reads using *Tosca* was very informative. Unique hybrid solutions were found for only 46.1% (6,835) of the reads (Suppl. Fig. S2A). Crucially, however, these solutions matched the known transcripts and coordinates in 97.2% of cases (Suppl. Fig. S2A). The main differentiating factor appeared to be the length of the sequencing read, with unrecovered reads significantly shorter (Suppl. Fig. S2B, median 58 v 68 nt, p < 2.2 × 10^−16^). Furthermore, the length of the shorter arm fragment within a hybrid read was also significantly shorter (Suppl. Fig S2C, median 21 nt v 28 nt, p < 2.2 × 10^−16^). It is likely that these shorter fragments align to multiple loci on the transcriptome and thus cannot be assigned uniquely; this prevents the reconstruction of a unique hybrid solution for these reads. There were no differences between high or low RNase digestion conditions (Suppl. Fig. S2B, C). Generally, a much smaller proportion of inter-transcript hybrids could be recovered (Suppl. Fig S2D), however there were unrecovered hybrid arms across all RNA types and regions (Suppl. Fig. S2E).

Importantly, this analysis using real (rather than synthetic) data shows that our computational approach can recover direct proximity ligation hybrids with high accuracy. It also demonstrates the importance of an adequately long cDNA to be able to do so. There is a balance between the level of RNase digestion, sequencing read length and maintaining the spatial resolution of the identified structure: we would recommend reads at least 75, if not 100 nt long and that RNase digestion is titrated so that each hybrid arm is at least 25-30 nt long.

### Non-hybrid reads help derive STAU1-bound short-range duplexes

While the new methods described thus far have considerably improved the recovery of hybrids from the STAU1 hiCLIP data, for 2,383,462 and 2,946,736 of the reads in the high and low RNase conditions we were ultimately unable to resolve a hybrid after mapping and hybrid identification and were thus non-hybrid reads. Owing to the stringent purification steps, these non-hybrid reads still represent sites where STAU1 has crosslinked to specific transcripts and thus reflect STAU1 binding. We set out to explore whether they could additionally represent short duplex structures whose loop regions were protected from RNase digestion (Fig. 1A, turquoise panel). We anticipated that such duplexes would have short loops and so to search for them, we developed a computational workflow to derive stem-loops near STAU1 crosslink peaks. We focused on 3’ UTRs because the majority of STAU1 crosslinking peaks (60.8%) are located in this region (Fig. 3A), but this approach is extendible to other transcript regions.

To explore the secondary structure around 3’ UTR STAU1 crosslink peaks, we first calculated the local base-pairing probability of each nucleotide and generated a meta-profile centred on the peak start position (Fig. 3B). As expected, there is a sharp trough in base-pairing probability around position 0, consistent with UV-crosslinking generally being much more efficient at unpaired nucleotides (25). Strikingly, there is an “M”-shaped profile in the +10 to +75 nt region (shaded grey), that reflects a tendency for a paired-unpaired-paired secondary structure motif downstream of the peak start, viz. a stem loop structure, with the paired regions the two arms of the stem, and a short intervening unpaired loop. This pattern is absent in control shuffled 3’ UTR sequences (dashed red line). Furthermore, this structural motif is absent around crosslink peaks of other known 3’ UTR binding single stranded RBPs, such as HuR and TDP-43 (Fig. 3B). Crosslinking between RBP and RNA primarily occurs at unpaired nucleotides, so STAU1 crosslinks are expected in three locations: upstream of a paired region, in the loop itself or downstream of a paired region. However, in practice, the experimental method and library preparation will advantage reads that contain at least part of the paired region as they are protected from the single-stranded RNase leaving longer RNA fragments (2). Furthermore, crosslinking is less likely to occur in the hairpin loops, given our prediction that they are likely to be short. Thus, STAU1 crosslinking upstream of a paired region is likely to dominate, as we observe here with the predicted “M” structural motif.

Next we focused on the individual peaks to characterise the presence of this structural motif further (Fig. 3C). We used k-means clustering to group peaks based on the pairing probability profiles in the +10 to +75 nt region. Applying the silhouette method identified two major clusters of non-hybrid peaks: those containing the “M”-shaped profile (7,567, 66.2%) and those that did not (3,861, Fig 3C, Cluster 0; Suppl. Fig. S3A). The peaks with a “M”-shaped profile were further grouped into 3 separate sub-clusters (Fig. 3C, Clusters 1-3), each differing in the distance from the STAU1 crosslink peak start (Cluster 1 closest starting at 30 nt, followed by Cluster 2 at 36 nt and Cluster 3 furthest away at 44 nt).

Finally, we predicted the RNA secondary structures to which these pairing probabilities correspond. For each cluster we defined the regions that are most likely to form duplexes by calculating the local minima of the pairing probability metaprofile (Suppl. Fig. S3B). This identified the duplex-containing regions as positions 1 to 30 (proximal arm) and 31 to 50 (distal arm) for Cluster 1; 8 to 36 and 37 to 63 for Cluster 2; and 16 to 44 and 45 to 68 for Cluster 3. Then we derived the corresponding RNA secondary structures by computationally hybridising the proximal and distal arm sequences (Fig. 3D). The majority of derived duplexes (93.6%) were 8 bp or longer with hybridisation energies indicating they were more stable than their shuffled controls (Fig. 3E, p < 2.2 × 10^−16^). NMR studies have found that stems of at least 8 bp were needed for STAU1 to bind (26), thus we excluded the small fraction (4%) of derived duplexes that were shorter than 8 bp from the subsequent analysis as they likely did not represent robustly bound structures (Fig. 3E). Thus, we obtained a final set of 6,995 duplexes that we term “derived” duplexes.

Our hypothesis was that these structures avoided digestion by RNase treatment owing to their short loops and steric protection by the bound STAU1 protein. Hence we assessed the loop lengths of our derived structures (Fig. 3F). The median loop length was indeed short at 7 nt with 75% of the loops less than 11 nt. Furthermore, we would expect these structures to be largely absent from the atlas of duplexes derived from hybrid reads.

We pooled all the experimentally-detected proximity ligation hybrids (linker and direct) and the derived duplexes, and applied our graph-based clustering approach to consolidate them into a complete atlas of STAU1-bound duplexes (Fig. 3G). We assessed how many duplexes were supported by one or more sources: 2,980 were detected by proximity ligation, of which 727 were supported by both direct ligation and linker hybrids. As expected, only a small fraction of the atlas (159; 1.6%) of duplexes overlapped derived structures (Fig. 3G). Altogether, using the graph-based clustering (rather than the less stringent original coverage-based approach) to identify experimentally-supported duplexes and incorporating the predicted duplexes, we recovered 9,766 STAU1-bound duplexes (excluding rRNA and tRNAs) (Fig. 3G), a ∼10-fold increase from the original.

### *Tosca*: a Nextflow pipeline for proximity ligation data analysis and visualisation

We developed a Nextflow computational pipeline, *Tosca*, to enable robust, reproducible and scalable implementation of our analysis approach (https://github.com/amchakra/tosca). Reference transcriptomes are currently provided for human and mouse, but custom references can also be user-generated. *Tosca* performs transcriptomic alignment and hybrid read identification and PCR de-duplication. This is followed by conversion to genomic coordinates and hybrid annotation. Graph-based clustering of hybrids enables delineation of the duplexes they represent. For these, hybridisation energies are calculated to assess duplex stability (Suppl. Fig. S1).

Importantly, *Tosca* also focuses on data visualisation and exploration. It generates BAM files that can be loaded in IGV as tracks to interactively display hybrids in the genome browser (Suppl. Fig. S4) with additional metadata enabling grouping and/or colouring by experiment or sample, duplex cluster, hybrid read orientation or hybridisation energy. Additionally, arc files for IGV can be generated for user-specified genes. Furthermore, files are generated to enable easy static plotting as arc plots (Fig. 4A) or as contact map matrices (Fig. 4B).

The *Tosca* pipeline can be run from the command-line using a single command to Nextflow that specifies the input files, transcriptome and additional customisable parameters. All the dependencies are containerised using Docker to ensure hassle-free deployment across platforms. We have provided user documentation in the repository and *Tosca* remains under active development.

### Comparison with global duplexes reveals insights into the RNA selectivity of STAU1

Our computationally-enhanced atlas finally enabled us to proceed with contextualising the STAU1-bound structures in the landscape of RNA structures globally. We used publicly available psoralen-based proximity ligation data in HEK293 cells generated using PARIS as a description of global RNA structures: both bound and not-bound by RBPs (4). To ensure a valid comparison, we re-processed the PARIS data using *Tosca* with the same gene annotations as for the hiCLIP analysis, and clustered the hybrid reads to generate an atlas of 494,174 duplexes (excluding rRNA and tRNA) from 14,745,302 unique hybrids across three biological replicates.

STAU1 duplexes recovered experimentally (i.e. through proximity ligation) likely reflected a separate group of STAU1-bound structures compared to those recovered through computational derivation (for which only 3’ UTR regions were considered) (Fig. 4A, B). We kept them separate for the subsequent comparative analyses to explore this further, and to avoid introducing bias to comparisons with PARIS. The regional distribution of duplexes for PARIS was similar to that originally obtained, with the majority in the CDS, followed by the 3’ UTR. This compared with the vast majority of STAU1-bound duplexes (81.3%) linking 3’ UTR regions (Fig. 5A). Given the strong selectivity of STAU1 for 3’ UTRs, we focused all subsequent comparisons on 3’ UTR intra-transcript duplexes: 2,368 were experimentally identified by STAU1 hiCLIP, and 28,306 by PARIS. Consistent with the original findings, there was only a small overlap between the two sets of duplexes (2.1%) (Fig. 5B) likely reflecting the low sensitivity of both these methods.

As previously noted, STAU1 often binds long-range 3’ UTRs duplexes, which can span intervening regions over 100 nt in length, and in some cases even kilobases (2). The proximity ligation atlas maintained the bimodal distribution of 3’ UTR duplex spanning loop lengths for STAU1 observed earlier (Fig. 2F, 5C). Fitting a two-component Gaussian mixture model revealed a primary peak (lambda = 0.60) with a mean duplex span of 233 nt and a secondary peak (lambda = 0.40) with a mean duplex span of 21 nt (Suppl. Fig. S5A). For global RNA duplexes from PARIS, however, there was a unimodal distribution with a median duplex span of 84 nt (Fig. 5C). Overall, this reinforces the finding that STAU1 preferentially binds long-range 3’ UTR duplexes, rather than this being a general feature of 3’ UTR duplexes.

Given the non-uniform distribution of duplex spans for STAU1-bound duplexes, we classified them into short-range (< 25 nt, 532 duplexes), medium-range (25-100, 677 duplexes), long-range (>100 nt, 1,132 duplexes) and derived (6,796 duplexes). Furthermore, we also found that while both STAU1 and PARIS duplexes were more thermodynamically stable than their shuffled controls (p < 2.2 × 10^−16^, and p < 2.2e × 10^−16^, Fig. 5D), STAU1 proximity ligation duplexes were also on average more stable than PARIS duplexes (p = 2.17 × 10^−145^) (Fig. 5D). While this could reflect a predilection of STAU1 to bind more stable structures, it is more likely to result from technical differences in the hiCLIP and PARIS experimental protocols. In hiCLIP, UV crosslinking of the RBP to the RNA is the only stabilising factor during the mild washing of beads performed before the proximity ligation, and so it is probable that less stable structures are lost through the lysis and washing steps. However, in PARIS psoralen directly crosslinks the RNA duplexes so they are more likely to be preserved through the experimental steps. Moreover, derived STAU1-bound duplexes have a similar hybridisation energy profile to PARIS duplexes, consistent with their representing weaker or transient interactions that are lost through the stringent hiCLIP washes.

Intriguingly, we noted that the hybridisation energies for the shuffled controls, where dinucleotide content had been preserved, were more negative for PARIS than for STAU1 hiCLIP. This led us to question whether there were differences in GC content; STAU1 hiCLIP duplexes were indeed significantly more AU-rich when compared with PARIS duplexes (Fig. 5E) and furthermore this was most dramatically the case for the long-range duplexes (46.7% v 33.3%, p = 5.97 × 10^−99^).

Examining the duplex structures in more detail, we observed that more of the residues in the STAU1 duplexes were involved in base-pairing than in the PARIS duplexes (Fig. 5F, median 84% v. 78%, p = 3.4 × 10^−164^). Furthermore, we found that the stem regions of 3’ UTR STAU1 proximity ligation duplexes with stems were on average longer than PARIS duplexes (median 21 v. 15, p < 2.2 × 10^−16^, Fig. 5G). This prompted us to explore the composition of the duplexes in terms of bulges (i.e. unpaired residues within the stem). We found that while there were similar numbers of bulges overall in the two (Suppl. Fig. S5B, median 3 v. 3 for proximity ligation duplexes, and 2 for predicted duplexes), their positioning differed: symmetry was an important distinguishing feature. While both sets had a small fraction of perfect duplexes and consisted mostly of asymmetric duplexes, for STAU1 hiCLIP, there was a higher proportion of symmetrical duplexes (i.e. with identically-sized bulges in both arms at the same position within the stem), 15.2-20% (mean: 16.8%) v. 6.4-8.1% (mean: 7.4%), despite these duplexes being longer (Fig. 5H). Interestingly, STAU1-bound derived duplexes share symmetry characteristics with proximity ligation duplexes (Fig. 5H), supporting that they are specific for STAU1, but thermodynamic features with PARIS duplexes (Fig. 5D), likely reflecting that they capture weaker or more transient interactions lost during the hiCLIP experimental steps.

Given the observed selectivity for symmetry in STAU1 hiCLIP and the preponderance of asymmetric duplexes, we characterised them in more detail. Even among these, STAU1 hiCLIP proximity ligation duplexes displayed a higher degree of symmetry, indicated by a higher percentage of all bulges having a symmetrically positioned bulge in the other arm (Suppl. Fig. S5C), whereas many asymmetric PARIS duplexes did not have a single position with symmetric bulges. Additionally, asymmetric STAU1 proximity ligation duplexes tend to have longer bulge-free stem segments (i.e. larger stretches with contiguous pairing) compared to PARIS (Suppl. Fig. S5D).

Although our observed STAU1 feature preferences may in part be attributable to technical differences between hiCLIP and PARIS (e.g. duplex stabilisation and purification methods), our findings are consistent with other biochemical and modelling studies. Intriguingly, the original NMR studies exploring the mechanisms of Staufen double-stranded RNA binding domain recognition of dsRNA found, through generation of synthetic stem loops that disruption of the helical structure through the introduction of bulges significantly reduced binding (26). Furthermore, our results are consistent with a previous computational model of *Drosophila* Staufen binding that found a preference for stems in which at least one side spans 12 nt and with few unpaired bases or internal loops (27). The preservation of structural symmetry may reflect the importance to STAU1 binding of a particular kind of tertiary conformation.

In summary, the principal features of STAU1-bound duplexes remain consistent across the span of duplexes, with an increase in A:U base pairs and symmetry for the long-range interactions spanning more than 100 nt. By comparing STAU1 hiCLIP detected duplexes with global 3’ UTR RNA duplexes determined using PARIS, we have been been able to assess characteristics of the proximity ligation duplexes that are enriched in the former, such as 3’ UTR binding, long-range loops, and symmetrical stems with more bases paired, all suggestive of the nature of STAU1 binding to RNA structures (Fig. 5I). The key features distinguishing the duplexes recovered by the two proximity ligation methods are highlighted in two illustrative examples on the *SRSF1* 3’ UTR (Fig. 5I, Suppl. Fig. S5E). Evidence to-date suggests that STAU1 likely recognises a predominantly structural, rather than sequence, motif through interactions with the dsRNA backbone (27, 28). Our findings support this hypothesis. However, while the GC-rich long base-paired regions of the well-studied *ARF1* binding site are thought to contribute to duplex stability, our data suggest a more complex picture related to the span of the given duplex. Notably, long-range duplexes are AU-rich, and therefore thermodynamically less favourable. However, they have longer, more symmetrical stems, with more base pairing (Fig. 5I) that is not evident in the global duplexes detected by PARIS. This requirement for specific lengths and structural conformations, rather than thermodynamic stability, to enable regulatory function is reminiscent of one recently described in the context of ribosome stalling (29).

### Integration with RNA metabolism reveals a stabilising role of selected 3’ UTR RNA duplexes

STAU1 is known to be involved in RNA metabolism, most notably RNA degradation (7, 8), but also translation (30). To explore the role of RNA structure in this regulation, specifically STAU1-bound duplexes, we integrated our expanded atlas with publicly available functional RNA data also in HEK293 cells that used metabolic labelling (4sU-seq) and mathematical modelling to measure synthesis, processing and degradation rates of RNAs (5). Given the predominance of STAU1 3’ UTR binding we again focused on this region.

First, we categorised genes into three groups on the basis of sources of their detected 3’ UTR duplexes: i) containing STAU1 (and/or PARIS) duplexes (STAU1 genes), ii) containing only PARIS duplexes (PARIS genes) or iii) not containing any detected duplexes. We then used the gene copy numbers calculated using spike-ins by (5) to assess for differences in RNA abundance (Suppl. Fig. S6A). We noted that genes in which no 3’ UTR duplexes were detected had a significantly lower expression level than those with PARIS or STAU1 duplexes. This marked difference likely reflects the sensitivity and detection limits of the methods. Consequently, for the subsequent comparative analysis we curated a set of 5,207 genes with an expression level greater than the 5th percentile of genes with PARIS or STAU1 duplexes.

We used the calculated RNA metabolism rates to assess the different levels of synthesis, processing and degradation for these two categories of genes with identified 3’ UTR duplexes (Fig. 6A). This showed STAU1 genes to have significantly higher synthesis and processing rates than genes without duplexes (Mann-Whitney test, p < 2.2 × 10^−16^ and p < 2.2 × 10^−16^ respectively) and significantly lower degradation rates (Mann-Whitney test, p < 3.4 × 10^−6^). There was also a similar pattern when compared to genes with PARIS duplexes (Suppl. Fig. S6B). Given the known role of STAU1 in RNA degradation through Staufen-mediated decay (7) this was a surprising finding. However, Staufen-mediated decay may only apply to a subset of transcripts. At a transcriptome-wide level, genes with STAU1-bound 3’ UTR duplexes have a lower degradation rate than genes with 3’ UTR duplexes without STAU1 binding. This prompted us to explore this relationship further.

Examining the distributions of RNA metabolism rates of the STAU1 genes revealed heterogeneity in their profiles (Fig. 6B). By using k-medoids clustering we could resolve the genes into three distinct clusters: A) with low synthesis and processing rates and high degradation rates; B) with intermediate synthesis, processing and degradation rates; and C) with high synthesis and processing rates and low degradation rates. Interestingly, cluster C - with the low degradation rate - also had a marked shift in the distribution of 3’ UTR duplexes span towards long-range duplexes (Fig. 6C). This was not reflected in the overall 3’ UTR length, where there was a significant shortening in 3’ UTR length with reducing degradation rates (Fig. 6D, Kruskal-Wallis test, p = 3.08 × 10^−10^) as expected given the known relationship between 3’ UTRs and RNA stability (31–33).

It appears that the long-range 3’ UTR duplexes bound by STAU1 “protect” against the degradation effect of longer 3’ UTRs. To quantify this we adapted the concept of circularisation score (11) to the 3’ UTR (the ratio of the STAU1 duplex span to the corresponding 3’ UTR length): a higher score indicates more compaction of the 3’ UTR by the STAU1 duplexes (Fig. 6E). This showed that cluster C had a significantly higher circularisation score than either A or B (Fig. 6E, Dunn’s test, p = 6.37 × 10^−6^ and p = 1.23 × 10^−5^ respectively).

To explore this effect in an alternative way, we assessed the positions of the proximal and distal duplex arms on the 3’ UTR (Fig. 6F). We divided 3’ UTRs into thirds and calculated which thirds were spanned by the two arms. This showed that duplexes generally tended to be located towards the start of 3’ UTR. Furthermore, longer-range duplexes spanning more thirds were proportionally greater in cluster C, again supporting a relationship between duplex span and degradation rates. Altogether, this suggests that there are two ends to the functional spectrum of genes with STAU1-bound duplexes (Fig. 6G). One (represented by cluster A) has short range structures, often at the proximal end of the 3’ UTR, reflected in a low circularisation score and with a high degradation rate; the other (represented by cluster C) has long-range duplexes that can span the whole 3’ UTR, compacting it, reflected in a high circularisation score, and with a lower degradation rate. Thus, we can link these RNA structures to RNA degradation, and suggest a potential protective effect of these long-range duplexes on degradation.

In conclusion, first, we have extended the atlas of STAU1-bound duplexes ∼10-fold primarily through demonstrating that i) direct proximity ligation occurs at sufficient levels in the absence of a linker adapter; and ii) short-range STAU1-bound duplexes exist, but cannot be recovered through proximity ligation and require computational derivation. Thus we propose that the linker adapter can be omitted from the hiCLIP protocol. Second, we have developed a robust, reproducible and scalable computational pipeline, *Tosca*, which is broadly applicable to multiple kinds of proximity ligation methods and includes downstream duplex characterisation and visualisation. Finally, we have integrated our atlas i) with global RNA duplexes to determine features of STAU1 RNA selectivity and ii) with RNA metabolism rates to uncover relationships between RNA structure and degradation.

## AVAILABILITY

All analysis code as well as RMarkdown notebooks to re-generate the figures is available at https://github.com/luslab/comp-hiclip. *Tosca* is available at https://github.com/amchakra/tosca.

## ACCESSION NUMBERS

Publicly available data was used for this project. Raw STAU1 hiCLIP data is available from ArrayExpress at E-MTAB-2937 with matched RNA-seq data at E-MTAB-2940. Raw PARIS data in HEK293 cells is available from GEO at GSE74353. Raw RNA metabolism data is available from GEO at GSE84722, with processed data obtained from the supplementary files. HuR and TDP-43 iCLIP data are available from ArrayExpress at E-MTAB-11854 and E-MTAB-4733 respectively.

## SUPPLEMENTARY DATA

Supplementary Data are available at NAR online.

## ACKNOWLEDGEMENTS

We would like to thank Neelanjan Mukherjee for sharing normalisation code for the RNA metabolism data. We would also like to thank the members of the Luscombe and Ule labs for fruitful discussions, in particular Dr Federico Agostini, Dr Charlotte Capitanchik and Dr Chris Cheshire.

