## Supplementary Figures for "A computationally-enhanced hiCLIP atlas reveals Staufen1 RNA binding features and links 3’ UTR structure to RNA metabolism"

### SUPPLEMENTARY FIGURE LEGENDS

**Suppl. Fig. S1: A workflow giving an overview of the main steps of the *Tosca* computational pipeline.**

**Suppl. Fig. S2: Analyses of in silico proximity ligation reads constructed from linker-containing hybrid reads.**
(A) Above: proportion of in silico reads recovered as hybrids using the Tosca pipeline. Below: the breakdown of recovered hybrids by whether they match the known true hybrid solutions.
(B) Read length distributions for recovered and not recovered hybrids across RNase conditions.
(C) Lengths of the shorter arms of hybrid reads for recovered and not recovered hybrids across RNase conditions.
(D) Proportions of inter-transcript and intra-transcript hybrids for recovered and not recovered hybrids
(E) Regional distribution of arms for recovered and not recovered hybrids

**Suppl. Fig. S3:**
(A) Silhouette method applied to the k-means clustering for Fig. 3C to identify the optimal number of clusters
(B) Mean pairing probability profiles for the 3 clusters identified in Fig. 3C. Dashed blue lines indicated local minima used to delineate the proximal and distal arm sequences.

**Suppl. Fig. S4: IGV screenshot of STAU1 hybrids on the ARF1, visualised using the BAM file and grouped by cluster using the "CL" tag and coloured by read orientation using the "RO" tag.**

**Suppl. Fig. S5:**
(A) Two-component Gaussian mixture model that identifies two duplex span size distributions for the STAU1 hiCLIP proximity ligation duplexes: red: $\mu = 22.75$, $\lambda = 0.41$; blue: $\mu = 242.95$ , $\lambda = 0.59$.
(B) Number of bulges per duplex for 3'UTR intra-transcript duplexes.
(C) Percentage of bulges with a symmetrically positioned and identical size bulge in the other arm within the duplexes.
(D) Longest bulge-free stem segment within the duplexes.
(E) Arc plot of the STAU1 hiCLIP and PARIS duplexes on the *SRSF1* 3' UTR, alongside STAU1 crosslinking signal and transcriptomic annotation; the duplexes used in Fig. 5I as representative for PARIS and STAU1 are indicated with a dagger and double dagger respectively.

**Suppl. Fig. S6:**
(A) RNA expression for genes with STAU1 3' UTR duplexes, global 3' UTR duplexes identified by PARIS (but not STAU1 hiCLIP) or without any identified 3' UTR duplexes.

(B) RNA metabolism (synthesis, processing and degradation rates) as measured by (5) for the genes as in (A), with those without any identified 3’ UTR duplexes subset to match the RNA expression of those with identified 3’ UTR duplexes either by STAU1 hiCLIP or PARIS.

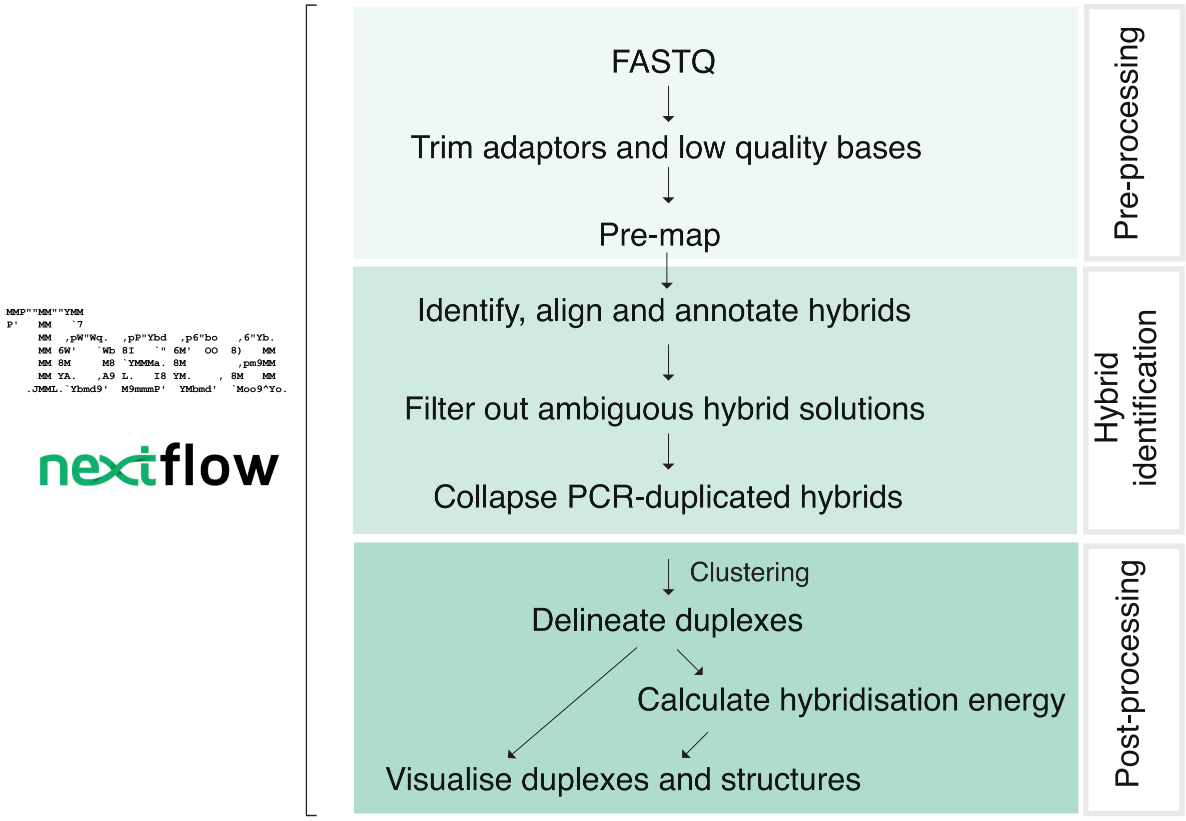
FASTQ
Trim adaptors and low quality bases
Pre-map
Pre-processing
Identify, align and annotate hybrids
Filter out ambiguous hybrid solutions
Collapse PCR-duplicated hybrids
Hybrid identification
Clustering
Delineate duplexes
Calculate hybridisation energy
Visualise duplexes and structures
Post-processing

**Suppl. Fig. S1**

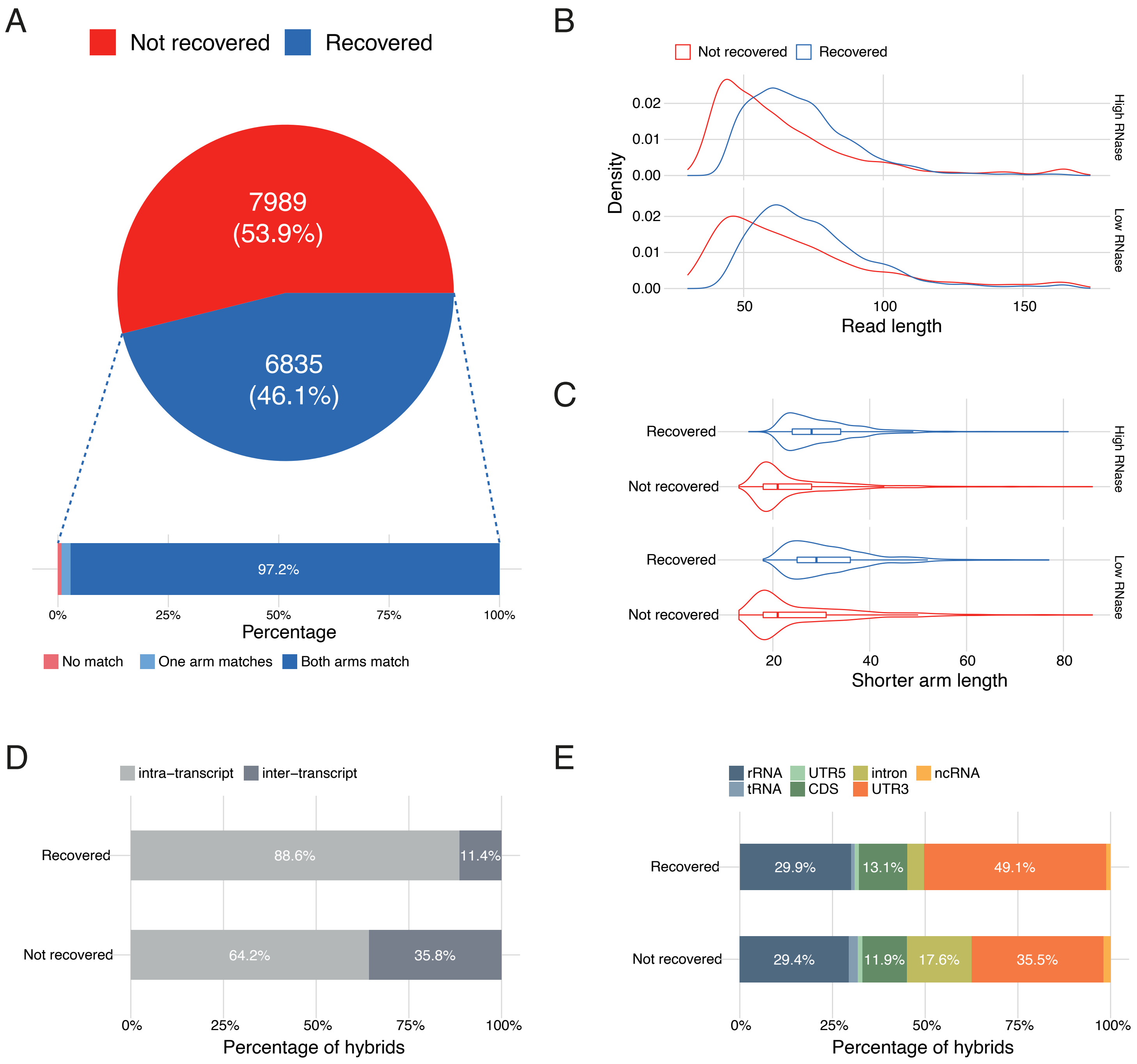
A
Not recovered
Recovered
7989
(53.9%)
6835
(46.1%)
97.2%
0%
25%
50%
75%
100%
Percentage
No match
One arm matches
Both arms match
B
Not recovered
Recovered
0.02
0.01
0.00
Density
High RNase
Low RNase
50
100
150
Read length
C
Recovered
Not recovered
High RNase
Low RNase
20
40
60
80
Shorter arm length
D
intra−transcript
inter−transcript
Recovered
Not recovered
88.6%
11.4%
64.2%
35.8%
Percentage of hybrids
E
rRNA
tRNA
UTR5
CDS
intron
UTR3
ncRNA
29.9%
13.1%
49.1%
29.4%
11.9%
17.6%
35.5%
Percentage of hybrids

**Suppl. Fig. S2**

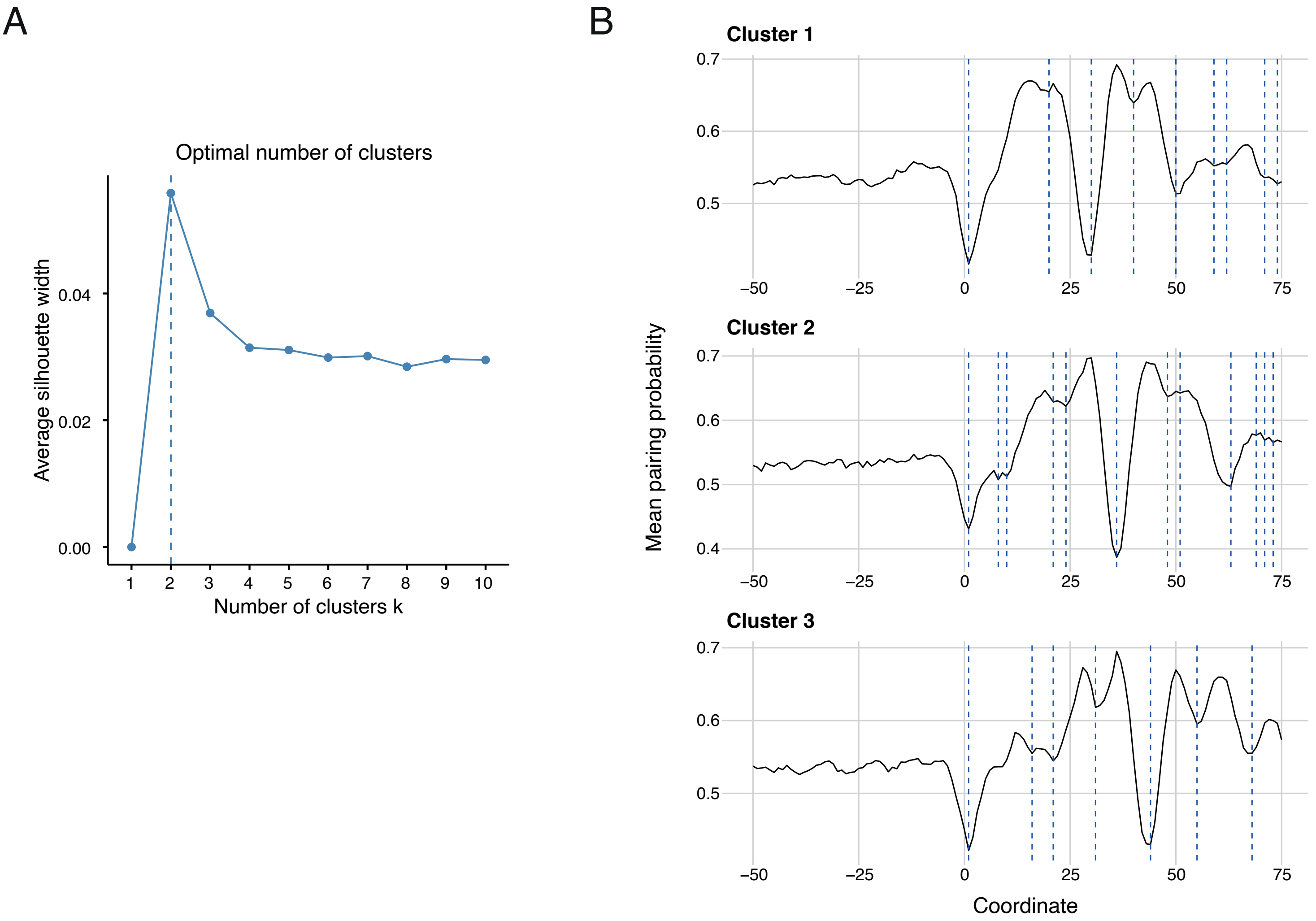
A
Optimal number of clusters
Average silhouette width
0.00
0.02
0.04
1
2
3
4
5
6
7
8
9
10
Number of clusters k
B
Cluster 1
Cluster 2
Cluster 3
Mean pairing probability
0.7
0.6
0.5
0.4
−50
−25
0
25
50
75
Coordinate

**Suppl. Fig. S3**

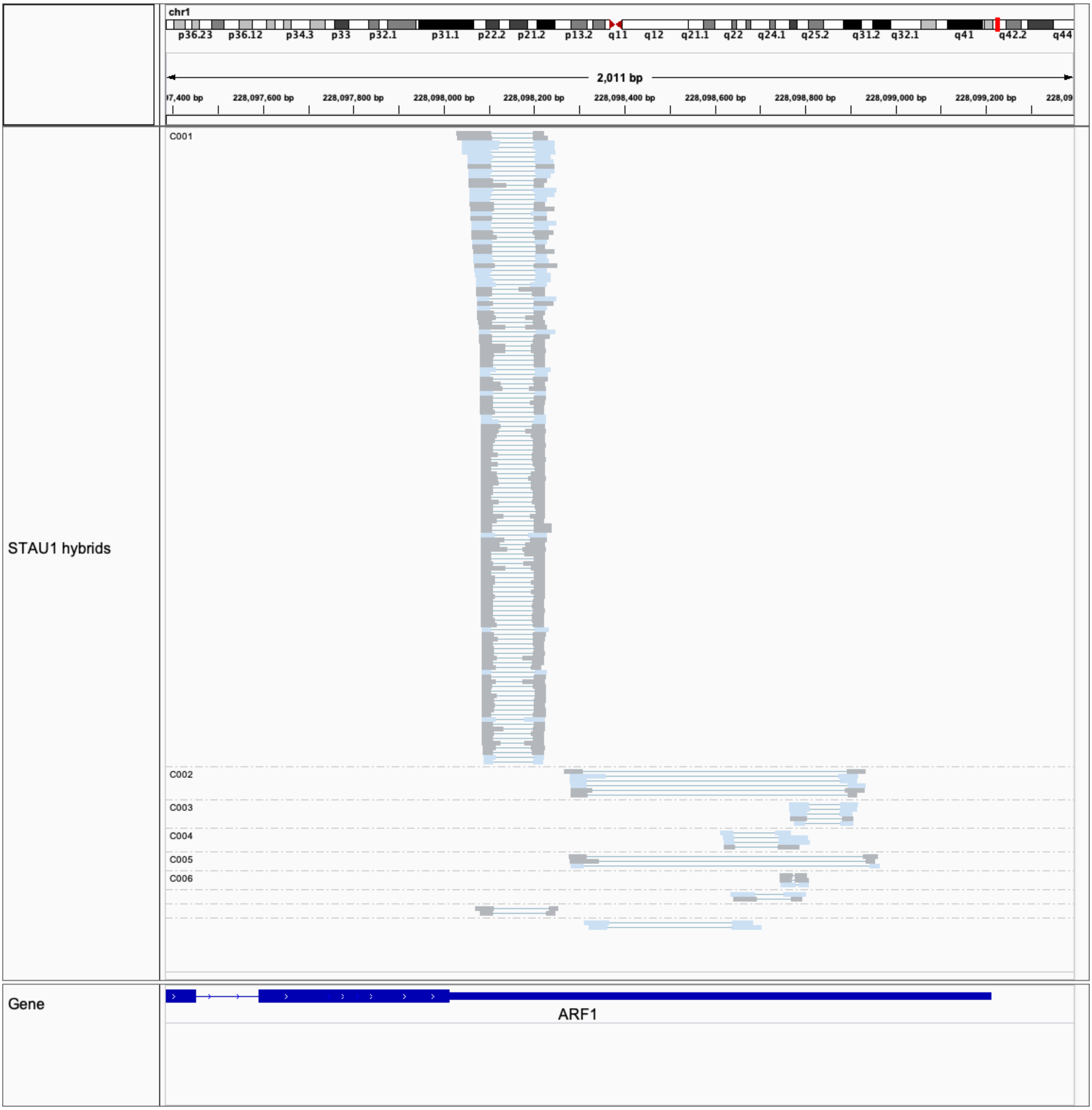

chr1
p36.23 p36.12 p34.3 p33 p32.1 p31.1 p22.2 p21.2 p13.2 q11 q12 q21.1 q22 q24.1 q25.2 q31.2 q32.1 q41 q42.2 q44
2,011 bp
228,097,600 bp
228,097,800 bp
228,098,000 bp
228,098,200 bp
228,098,400 bp
228,098,600 bp
228,098,800 bp
228,099,000 bp
228,099,200 bp
C001
C002
C003
C004
C005
C006
STAU1 hybrids
Gene
ARF1


**Suppl. Fig. S4**

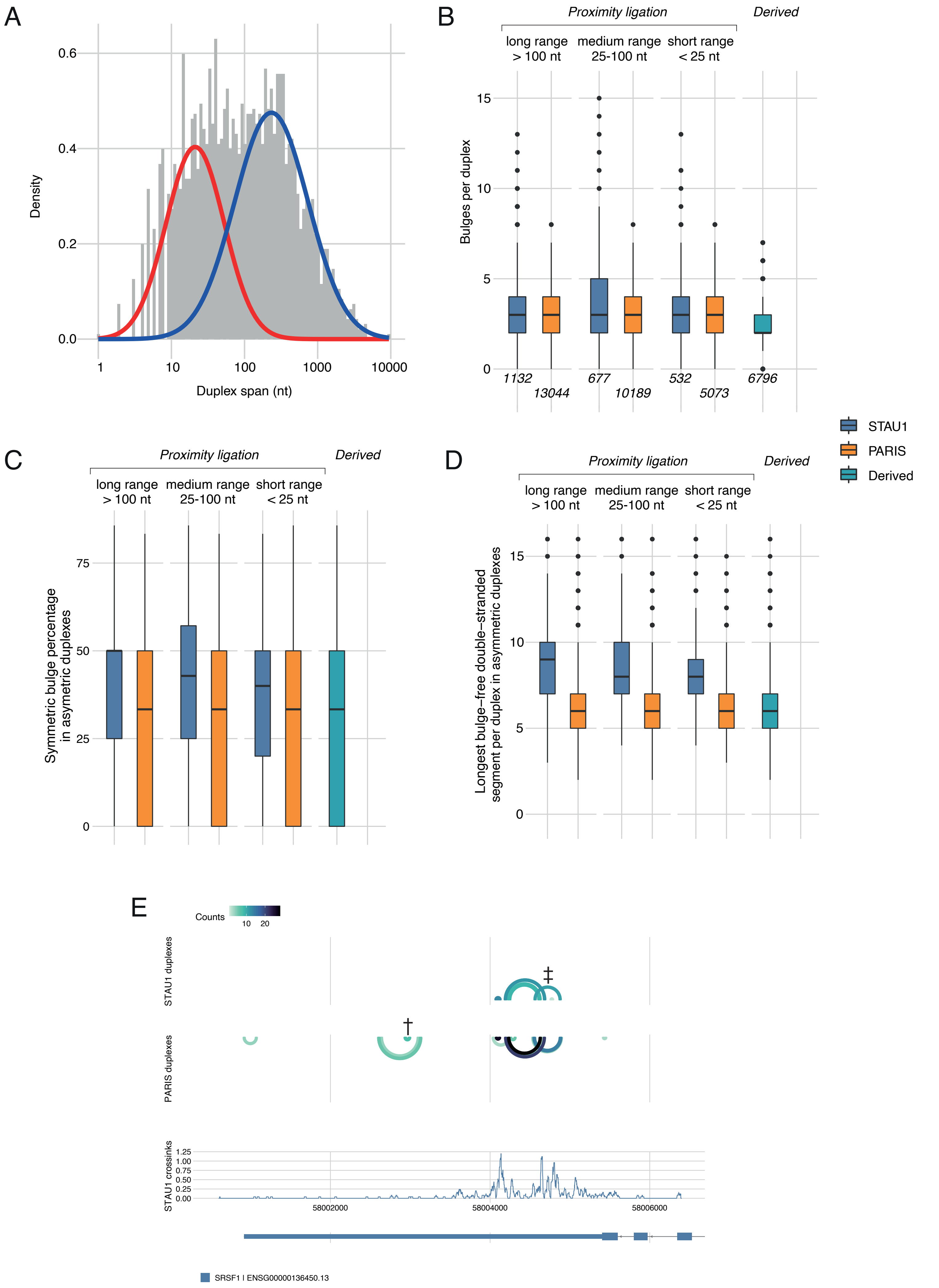
A
Density
Duplex span (nt)
B
Proximity ligation
Derived
long range > 100 nt
medium range 25-100 nt
short range < 25 nt
Bulges per duplex
1132
13044
677
10189
532
5073
6796
STAU1
PARIS
Derived
C
Symmetric bulge percentage in asymmetric duplexes
D
Longest bulge−free double−stranded segment per duplex in asymmetric duplexes
E
Counts
STAU1 duplexes
PARIS duplexes
STAU1 crossinks
58002000
58004000
58006000
SRSF1 | ENSG00000136450.13

**Suppl. Fig. S5**

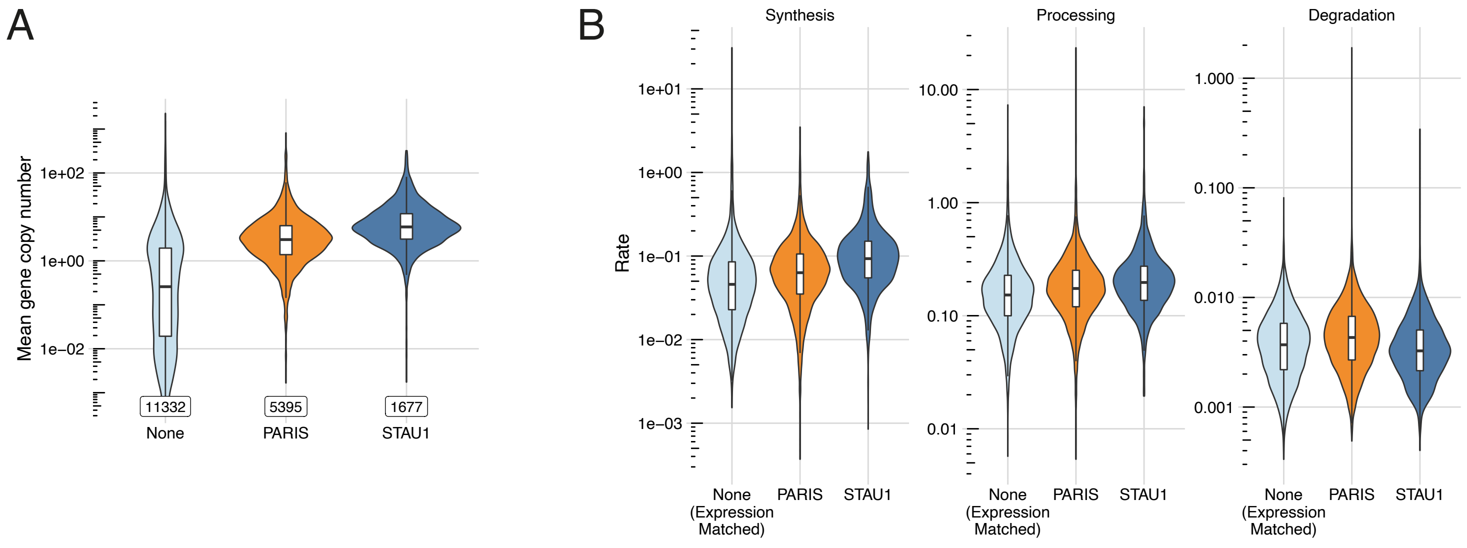
A
B
Mean gene copy number
1e+02
1e+00
1e−02
11332
5395
1677
None
PARIS
STAU1
Synthesis
Processing
Degradation
Rate
1e+01
1e+00
1e−01
1e−02
1e−03
10.00
1.00
0.10
0.01
1.000
0.100
0.010
0.001
None (Expression Matched)
PARIS
STAU1

**Suppl. Fig. S6**
